# Neutrophil counts and cancer prognosis: an umbrella review of systematic reviews and meta-analyses of observational studies

**DOI:** 10.1101/330076

**Authors:** Meghan A. Cupp, Margarita Cariolou, Ioanna Tzoulaki, Evangelou Evangelos, Antonio J. Berlanga-Taylor

## Abstract

**OBJECTIVE:** To evaluate the strength and validity of evidence on the association between the neutrophil to lymphocyte ratio (NLR) or tumour-associated neutrophils (TAN) and cancer prognosis.

**DESIGN:** Umbrella review of systematic reviews and meta-analyses of observational studies.

**DATA SOURCES:** Medline, EMBASE, and Cochrane Database of Systematic Reviews.

**ELIGIBILITY CRITERIA:** Systematic reviews or meta-analyses of observational studies evaluating the association between NLR or TAN and specific cancer outcomes related to disease progression or survival.

**DATA SYNTHESIS:** The available evidence was graded as strong, highly suggestive, suggestive, or weak through the application of pre-set grading criteria. For each included meta-analysis, the grading criteria considered the significance of the random effects estimate, the significance of the largest included study, the number of studies and individuals included, the heterogeneity between included studies, the 95% prediction intervals, presence of small study effects, excess significance and credibility ceilings.

**RESULTS:** 239 meta-analyses investigating the association between NLR or TAN and cancer outcomes were identified from 57 published studies meeting the eligibility criteria, with 81 meta-analyses from 36 studies meeting the criteria for inclusion. No meta-analyses found a hazard ratio (HR) in the opposite direction of effect (HR<1). When assessed for significance and bias related to heterogeneity and small study effects, only three (4%) associations between NLR and outcomes in gastrointestinal and nasopharyngeal cancers were supported by strong evidence.

**CONCLUSION:** Despite many publications exploring the association between NLR and cancer prognosis, the evidence is limited by significant heterogeneity and small study effects. There is a lack of evidence on the association between TAN and cancer prognosis, with all nine associations identified arising from the same study. Further research is required to provide strong evidence for associations between both TAN and NLR and poor cancer prognosis.

**REGISTRATION:** This umbrella review is registered on PROSPERO (CRD42017069131)

**FUNDING:** Medical Research Council

**COPYRIGHT:** Open access article under terms of CC BY

**SHORT TITLE:** Neutrophils and cancer prognosis: an umbrella review

**KEY RESULT:** When assessed for significance and bias related to heterogeneity and small study effects, only three (4%) associations between NLR and overall survival and progression-free survival in gastrointestinal and nasopharyngeal cancers were supported by strong evidence.

**WHAT THIS PAPER ADDS:** 

**WHAT IS ALREADY KNOWN ON THE TOPIC:**

- Neutrophil counts have been linked to the progression of cancer due to their tumourigenic role in the cancer microenvironment.
- Numerous meta-analyses and individual studies have explored the association between neutrophil counts and cancer outcomes for a variety of cancer sites, leading to a large body of evidence with variable strength and validity.
- Uncertainty exists around the association between neutrophils and cancer outcomes, depending on the site, outcome and treatments considered.

**WHAT THIS STUDY ADDS:**

- All meta-analyses included in this review indicated an association between high neutrophil counts and poor cancer prognosis.
- There is strong evidence supporting the association between the neutrophil to lymphocyte ratio and poor cancer prognosis in some respiratory and gastrointestinal cancers.
- Further research is required to strengthen the existing body of evidence, particularly for the association between tumour-associated neutrophils and cancer outcomes.

## INTRODUCTION

Cancer is the second leading cause of death worldwide(1), contributing to over 8.7 million deaths globally(2). Cancer incidence is increasing(2), due in part to the epidemiological transition of increasing mortality and morbidity from chronic diseases(3). This increase highlights the importance of identifying prognostic indicators associated with cancer progression(4). C-reactive protein (CRP)(5), serum albumin(6), fibrinogen and differential leukocyte counts(7–9) are all indicators of inflammation that have been linked to cancer prognosis. In recent years, the role of neutrophils in the tumour microenvironment has been explored due to their paradoxical role in both the prevention and facilitation of tumour progression(10).

Neutrophils are the most abundant white blood cells (WBCs)(11), making up 50-70% of the body’s circulating leukocytes(12). Neutrophil counts, particularly the neutrophil to lymphocyte ratio (NLR), have emerged as indicators of cancer prognosis and several systematic reviews and meta-analyses have explored their potential as a prognostic indicator in cancer(13). The NLR was first recognised for its association with systemic inflammation in the critically ill(14) and meta-analyses on the association between elevated NLR and poor prognosis have reported a wide range of effect sizes depending on the site of cancer(13). It is currently unclear how the association between NLR and poor prognosis varies depending on the site of cancer or the treatment considered. The close association between inflammation and cancer progression indicates that elevated tumour-associated neutrophils (TAN), also known as neutrophils which infiltrate tumours(15), are a potential prognostic indicator(10,16,17). Many systematic reviews and meta-analyses explore the association between neutrophils and cancer prognosis. However, the myriad of different cancer sites, stages, treatments and survival outcomes measured complicates the interpretation of this body of evidence.

Umbrella reviews allow for the analysis of broad subject areas to examine the strength and credibility of associations using the results of published systematic reviews and meta-analyses(18,19). Through umbrella review methods, the strength and consistency of the literature is assessed to evaluate bias and identify which associations are supported by strong evidence(18). Here we carried out an umbrella review of systematic reviews and meta-analyses with the aim of comprehensively evaluating the validity and strength of reported associations between NLR or TAN and cancer prognosis and identify potential biases in relevant literature.

## METHODS

### Literature search

Searches were conducted in Medline, Embase and the Cochrane Database (*Appendix 1*) and aimed to include all systematic reviews and meta-analyses published in English from inception up to 23 June 2017. Indicators of neutrophil counts included NLR and TAN (intratumoural neutrophils, peritumoural neutrophils and stromal neutrophils). Overall survival (OS), cancer-specific survival (CSS), progression-free survival (PFS), disease-free survival (DFS) and reoccurrence-free survival (RFS) were considered as cancer outcomes. Articles were initially screened by title and abstract to determine eligibility for full text screening and inclusion using RefWorks web-based bibliography and database manager(20).

### Inclusion and exclusion criteria

Included studies were systematic reviews and meta-analyses of individual observational studies in humans with any cancer diagnosis and NLR or TAN measurements taken around the time of diagnosis. Systematic reviews which did not include a meta-analysis were excluded. Meta-analyses were excluded if they did not assess a cancer outcome in our inclusion criteria, included more than one outcome in a single analysis, or either did not specify the cancer site studied or included all sites in a single analysis (e.g. analyses of cancers grouped as “other” were excluded). Meta-analyses were also excluded if they did not provide sufficient detail for replication, such as the individual hazard ratio (HR), 95% confidence interval and total sample size of each included study. If a single study included multiple meta-analyses, all meta-analyses were individually assessed for eligibility.

When more than one meta-analysis was identified for a single association at a specific site they were assessed for concordance in the direction, magnitude and significance of their effect estimates. If the duplicate meta-analyses agreed in significance, magnitude and direction of effect, the meta-analysis with the greatest number of studies was included. If the duplicates had any disagreement, all meta-analyses were excluded for the association.

### Data extraction

Data extraction forms were generated to record information from each meta-analysis and the included individual studies. From each meta-analysis the study’s first author, year of publication, outcome measure, indicator and cancer diagnosis were extracted. For each included individual study, the first author, year of publication, total population, epidemiological design, HR and 95% confidence interval were extracted. Each meta-analysis was allocated to one of six categories according to cancer site as follows: all cancers, gastrointestinal, gynaecological, hepatocellular, respiratory and urinary cancers.

### Data analysis

#### Estimation of summary effects

The weighted inverse variance method was used to reproduce all included meta-analyses in R(21) with the “meta” package(22) and “metagen” command. For each cancer site specific indicator and outcome pair, the summary effect size and 95% confidence interval were calculated using fixed and random effects methods. The random effects model was used to compute summary effect size estimates taking into account the observed heterogeneity, since cancer is a highly heterogeneous disease(23,24). Estimates from the fixed effects model are also presented.

#### Assessment of reproducibility

Each included meta-analysis was reproduced to yield both fixed and random effects estimates. Reproduced random or fixed effect estimates which did not match the results reported in the original study were assessed for absolute and percent difference. Meta-analyses with a difference in HR of only 0.01 were attributed to rounding errors. Studies with larger discrepancies were investigated to determine the source of disagreement. Where there were issues with reproducibility, the calculated values of the random effects model were used to assess the evidence for the association.

#### Assessment of heterogeneity

Heterogeneity within each meta-analysis was assessed with Cochrane’s Q test and quantified using the I^2^ statistic(25). Cochrane’s Q test detects a departure from homogeny in the effect sizes of individual studies when p<0.10(25). The I^2^ statistic was also used to quantify the percentage of variation which can be attributed to heterogeneity due to common limitations associated with Cochrane’s Q test(25). Values exceeding 50% or 75% are considered to show large or very large heterogeneity respectively. The 95% confidence intervals around each I^2^ value were also included to evaluate the uncertainty around estimates of heterogeneity(26). Large measures of heterogeneity, representing true heterogeneity or inconsistency due to bias(27), were further assessed through prediction intervals and Egger’s test for funnel plot asymmetry.

#### Estimation of prediction intervals

In order to assess the impact of heterogeneity, 95% prediction intervals were calculated for the summary random effect estimates(28). Prediction intervals account for the uncertainty caused by heterogeneity when estimating the distribution of true effect sizes in an association and yield an interval which predicts the effect size of future studies investigating the same association(28). In studies with large amounts of heterogeneity the prediction interval may be wide enough to include the null value (HR<1), suggesting that the true effect size may also include the null value.

#### Assessment of small study effects

Small study effects and funnel plot asymmetry were quantified through Egger’s test using the command “metabias” from the R package “meta”(22) to determine if heterogeneity occurred due to chance(29). The presence of small study effects was confirmed by a low significance value in Egger’s test (p<0.10) indicating bias or true heterogeneity(30). Since the Egger’s test is underpowered in meta-analyses including less than ten individual studies(31), further assessment was carried out in these meta-analyses to determine if the summary effect size estimate was greater than the point estimate of the largest included study, indicating potential small study effects(32).

#### Evaluation of excess significance

The test for excess significance (TES)(33) was used to determine if the number of observed positive results differed significantly from the expected number of significant results. TES results can reveal reporting bias if the number of observed studies with significant results in each meta-analysis is significantly larger than the expected number using a two-tailed binomial probability test (p<0.10)(34). The expected number of significant results in each meta-analysis was calculated as the sum of the statistical power estimate, or the probability that each study will find a positive result(33,34). The estimated power for each individual study was calculated in Stata 14(35), using the “power cox” command to calculate the power of each test given its sample size, effect size and significance level(36). The estimation of power for each individual study also requires an estimation of the true effect size, so the effect size of the largest study was used to give the most conservative estimation of true effect. Estimates from both fixed and random effects models were included for sensitivity analysis. The “binom.test” command in R was used to assess the significance of differences in the number of observed versus expected significant studies through an exact binomial test(37).

#### Credibility ceilings

Credibility ceilings were utilised for sensitivity analysis and to test the methodological limitations of using observational studies to calculate combined effect estimates(38,39). Credibility ceiling calculations inflate the variance of each study included in a meta-analysis to account for the probability *c* that the true effect size is in the opposite direction of effect of the observed point estimate(39). Inflated variances were calculated in Stata 14(35,38). The summary effect size and heterogeneity of each meta-analysis was assessed with ceiling values ranging from 5 to 20%.

#### Grading the evidence

Associations between neutrophil counts and cancer prognosis were categorised into strong, highly suggestive, suggestive, or weak through assessment of the strength and validity of the evidence for each meta-analysis, according to pre-defined criteria outlined in *Supplementary Figure 1*(40,41). In order for an association to be considered strong, the meta-analysis must yield a p-value of less than 10^−6^ in the random effects model(42), include more than 1,000 individuals, show significance at p<0.05 in the largest included study, find no heterogeneity (p>0.10) through the Q test, detect less than 50% variance due to heterogeneity through the I^2^ statistic, yield a prediction interval excluding the null value (HR=1), display no evidence of small study effects or excess significance, and the association must maintain significance at p<0.05 with the application of a credibility ceiling of 10%. The number of studies in each meta-analysis was also included as eligibility criterion for strong evidence since a sample size greater than three is required for reliable assessment of heterogeneity and small study effects(25,31,43).

#### Quality assessment

Studies with meta-analyses categorised as providing either highly suggestive or strong evidence underwent quality assessment through AMSTAR 2, a tool for assessing the methodological quality of systematic reviews (44). Studies were assessed by two reviewers (MAC and MC) and consensus reached on any disagreements in quality. Statistical analyses were carried out R(21), including the packages “meta”(22) version 4.8-4 and “ggplot2”(45) version 2.2.1, and Stata 14(35).

### Patient involvement

No patients were involved in development of the study design nor were they asked to advise on interpretation. No ethical approval was required for this review since it relied entirely on anonymised, published data.

## RESULTS

### Characteristics of included meta-analyses

The 57 published articles meeting the criteria for inclusion contained 239 meta-analyses (*Appendix 2*). The 81 meta-analyses meeting the eligibility criteria arose from 36 of these articles, published between 2014 and 2017 (*Figure 1*)(46–81). These meta-analyses included individual studies which presented NLR or TAN categorically as either high or low and investigated 40 associations for 27 different cancer diagnoses, including nine subtypes for treatment and four for cancer stage (*Supplementary Table 1*). The meta-analyses were grouped as all cancers (n=8), gynaecological (n=6), gastrointestinal (n=24), hepatocellular (n=11), respiratory (n=10) and urinary cancers (n=22) (*Figures 2A and 2B*). Included meta-analyses summarised effect size estimates from 693 individual studies, with OS as the most frequently assessed outcome (n=41). In 51 meta-analyses (63%) total sample size exceeded 1,000 individuals and each meta-analysis had a median of five studies. However, 57 meta-analyses (70%) included less than ten studies and 17 (21%) included only two studies. The characteristics of included meta-analyses are summarised in *Supplementary Table 2*.

**Figure 1.**
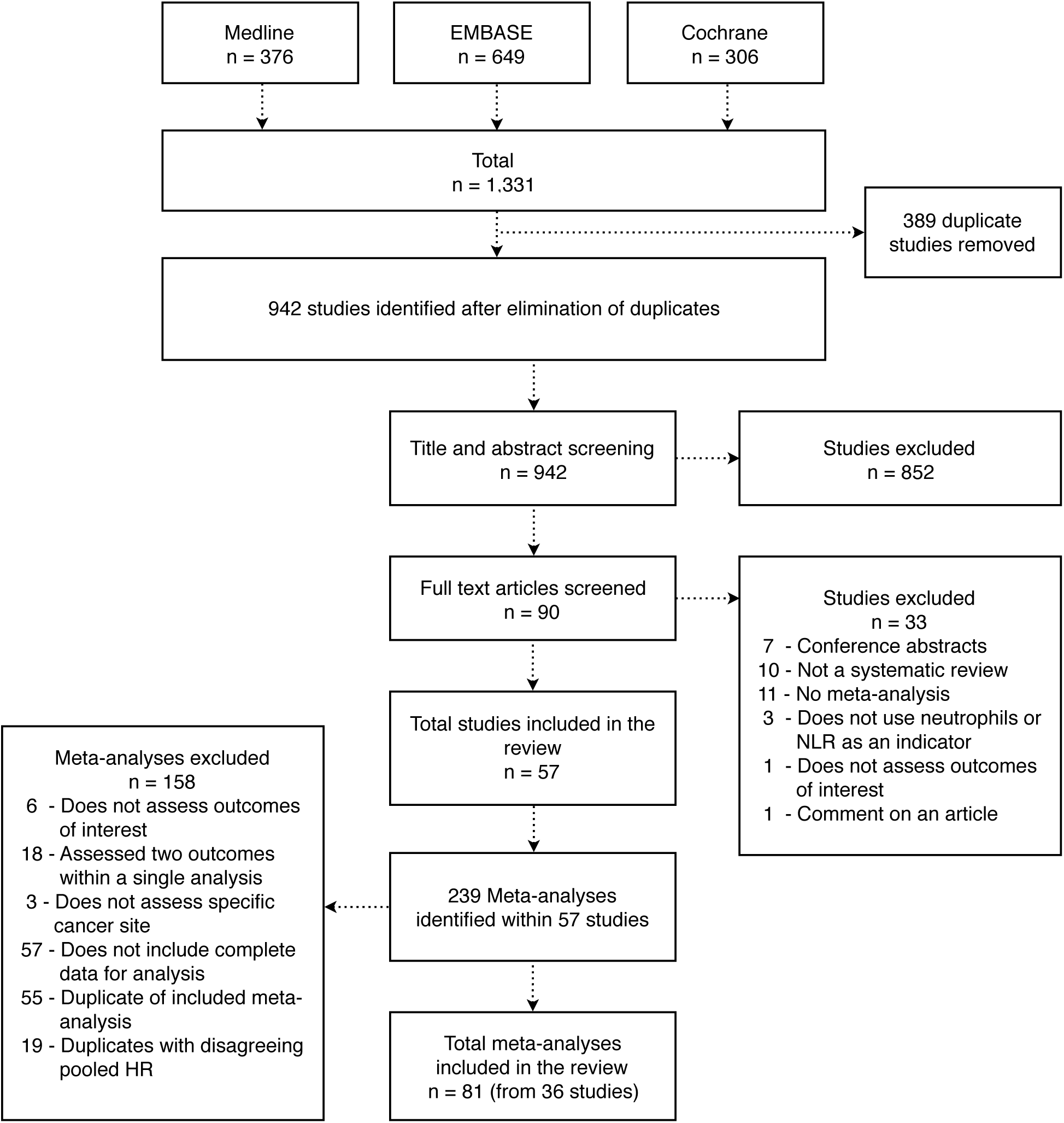
Flowchart of study and meta-analysis selection.

**Figure 2.**
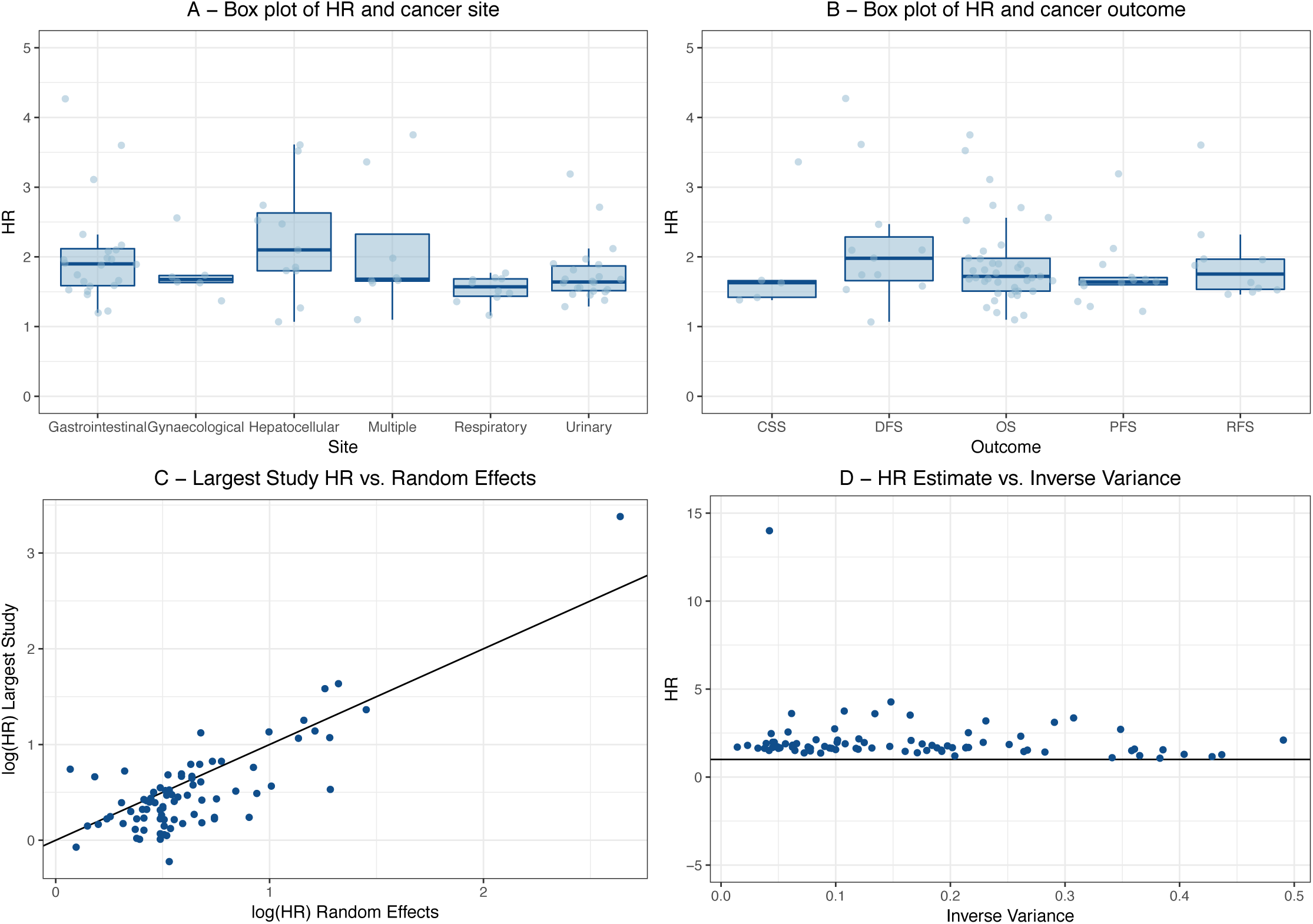
Assessment of consistency in meta-analyses. **A, B** - Box plots of random effects HR estimates for each meta-analysis by cancer site and outcome. The Y-axis labelled “HR” details the effect size for each meta-analysis describing an association between NLR or TAN and cancer prognosis for each site grouping. The X-axis labelled “Site” in Figure 2 A represents each site group meta-analyses have been sorted into. The multiple subgroup contains all cancers, defined as a grouping of cancer diagnosis unrelated to site, stage or treatment. The X-axis labelled “Outcome” in Figure 2 B represents the prognostic outcome assessed in each meta-analysis. The outlier of HR=14 for NLR and OS in rectal cancer has been excluded from these figures. **C** - Log(HR) of largest study versus log(HR) of random effects estimates for each meta-analysis. The Y-axis labelled “log(HR) Largest Study” represents the log of the HR of the largest study included in each analysis. The X-axis labelled “log(HR) Random Effects” represents the log of the HR of the random effects estimate calculated in each meta-analysis. **D** - Random effects estimates versus inverse variance. The Y-axis labelled “Random Effects Estimates Hazard Ratio” represents the HR of random effects estimate for each meta-analysis. The X-axis labelled “Inverse Variance” represents the inverse of the variance 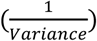 for each meta-analysis.

A total of 74 duplicate meta-analyses were excluded. Nineteen meta-analyses assessing six associations were excluded due to disagreement in significance between duplicates. A further 55 duplicate meta-analyses that agreed in significance, magnitude and direction of effect were excluded for 31 associations and only the meta-analysis with the largest number of studies was included for each association (*Appendix 3*).

### Summary effect size

All estimated summary effect sizes for both fixed and random effects estimates are shown in *Supplementary Figures 2-82*.

Using a threshold of p<0.05 for statistical significance, 72 of the 81 meta-analyses (89%) were significant with random effects. At a more stringent threshold of p<10^−6^, the number of statistically significant meta-analyses for random effects dropped to 38 (47%) (*Supplementary Table 2*). The 38 meta-analyses with significance at p<10^−6^ assessed both NLR and intratumoural neutrophils as indicators of poor prognosis in 19 cancer sites. Thirty-five of these 38 meta-analyses assessed NLR as an indicator of poor prognosis in gynaecologic, gastrointestinal, hepatocellular, respiratory, urinary, and all cancers. Intratumoural neutrophils were assessed as an indicator of poor prognosis in three of the 38 meta-analyses (8%), including urinary and all cancers.

In 20 meta-analyses (25%), the largest study included was not statistically significant at p<0.05. However, 16 of these meta-analyses still had a statistically significant summary random effects estimate. In two meta-analyses, the largest study included had an effect size in the opposite direction to the random effects estimate (HR<1). The largest study effect sizes tended to be more conservative estimates of effect than the random effect estimates, with 58 meta-analyses (72%) yielding a HR which was greater than the point estimate of the largest included study. However, there was correlation between the log(HR) of the summary random effects and the largest study for each meta-analysis, indicating consistency in the results (*Figure 2C*).

In order to determine the impact of study size on the magnitude of the summary effect size, random effects estimates were plotted against inverse variance for each meta-analysis. When compared to meta-analyses with large variances those with smaller variances studies produced more conservative estimates, displaying a smaller range of HR estimates and a slight tendency toward a null value (HR=1). Meta-analyses with large variance displayed a wide range of random effects HR and included an increased number of HR estimates greater than two (*Figure 2D*).

### Reproducibility

In 21 of the 81 included meta-analyses, the HR was reproduced imperfectly. Nine of the 21 imperfectly reproduced meta-analyses were within 0.01 of the reported HR, and the differences were attributed to rounding errors. The remaining 12 meta-analyses were within 10% of the reported HR, with six meta-analyses (50%) reporting an HR with less than a 2% difference from the calculated HR (*Appendix 4*).

### Heterogeneity between studies

Cochrane’s Q test was significant at p<0.10 in 40 of the 81 included meta-analyses (50%). In 20 meta-analyses (25%), the I^2^ statistic was greater than 75%, indicating a high level of variability between studies due to heterogeneity. An additional 19 meta-analysis yielded I^2^ values between 50% and 75% *(Appendix 5)*. In 30 meta-analyses (37%), the I^2^ statistic was less than 50%, but only six (9%) yielded a I^2^ statistic with a 95% CI that did not include 50%. The confidence interval of the I^2^ statistic was not used as criteria for grading evidence, since large fluctuations in I^2^ occur when meta-analyses include less than 15 studies(43).

Prediction intervals were not calculated for 17 (21%) meta-analyses which had included only two individual studies. The prediction intervals of 46 meta-analyses (57%) included the null value of HR=1. Of 64 meta-analyses (79%) including at least three individual studies, 18 had prediction intervals which excluded the null value (HR=1). The 12 meta-analyses (15%) including exactly three individual studies yielded very wide prediction intervals, all of which included the null value of HR=1.

### Small study effects

Sixty-five (80%) of the 81 included meta-analyses were judged to have evidence of small study effects *(Appendix 5).* Sixty-four meta-analyses included three or more studies and were eligible for further assessment through Egger’s test for asymmetry(22). Thirty-seven (58%) of these 64 meta-analyses yielded significant p-values (p<0.10). However only 24 (30%) meta-analyses included ten or more individual studies, a cut off required to give Egger’s sufficient power(31,32). Twenty-one (88%) of the 24 meta-analyses including 10 of more individual studies yielded a significant Egger’s test (79%), indicating funnel plot asymmetry(30).

Forty meta-analyses analysed between three and nine studies and Egger’s test was significant (p<0.1) in 16 of these (40%). In 11 of the remaining 24 meta-analyses (46%), the summary effects estimate from the random effects model was larger than the point estimate of the largest study and they were considered to have evidence of small study effects.

### Excess significance

Seventeen meta-analyses (21%) showed evidence of excess significance bias according to the TES when the effect size of the largest included study was utilised as an estimate of true effect size (*Appendix 6*). When the fixed summary effect sizes were utilised as an estimation of true effect size, fifteen meta-analyses (16%) showed evidence of excess significance. No meta-analyses showed evidence when the random summary effect sizes were used.

### Credibility ceilings

The summary effect size estimates and significance of each meta-analysis matched that of the random effects model at a credibility ceiling of 0%, with 72 of the 81 meta-analyses being significant at p<0.05 (89%) (*Table 1*). At a ceiling of 5%, 65 maintained significance (71%) and 50 (61%), 37 (46%), and 22 (27%) maintained significance at ceilings of 10%, 15%, and 20%, respectively. All of the meta-analyses remained consistent in direction of effect (HR>1) up to a ceiling of 15% and 2 (2%) yielded an effect estimate in the opposite direction (HR<1) with a ceiling of 20%. The I^2^ value of each meta-analysis decreased with each increase in ceiling value.

**Table 1.**
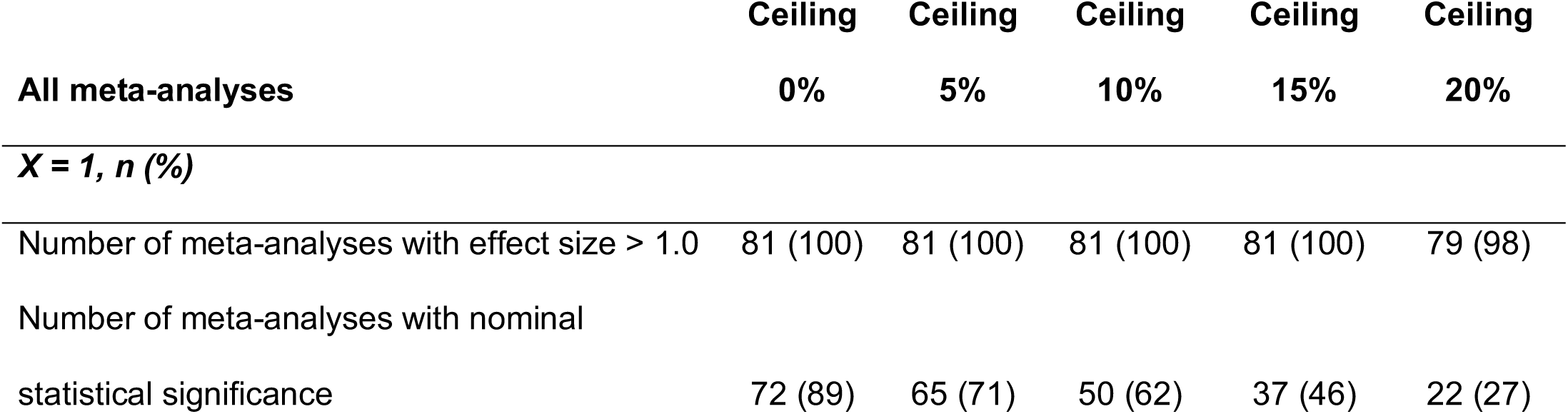
Credibility ceiling results

### Grading the evidence

Each included meta-analysis was evaluated to determine if the associations they explored was supported by strong, highly suggestive, suggestive or weak. In nine meta-analyses (11%), no significance was detected at a threshold of p<0.05. The remaining seventy-two meta-analyses (89%) provided at least weak evidence of an association (p<0.05) (*Table 2*).

**Table 2.** Grading of evidence

| Evidence | Criteria | Increased risk of poor prognosis |
| --- | --- | --- |
| Strong (n=4) | p<10<sup>-6</sup>* with random effects;<br>>1000 individuals included; >3 studies included; largest study significant at p<0.05; Q test significant at p<0.10; I<sup>2</sup> less than 50%, prediction interval does not include null value (HR=1); small study effects not detected; excess significance not detected | CLM-OS (NLR), CLM SR-OS (NLR), Nasopharyngeal-PFS (NLR) |
| Highly Suggestive (n=22) | p<10<sup>-6</sup>* with random effects;<br>>1000 individuals included;<br>largest study significant at p<0.05 | All cancers-DFS (NLR), All cancers-PFS (NLR), Breast-DFS (NLR), Breast-OS (NLR), Cervical-OS (NLR), Gynaecologic-OS (NLR), Gastric-OS (NLR), Pancreatic-OS (NLR), Colorectal-OS (NLR), CLM-RFS (NLR), HCC-OS (NLR), HCC-DFS (NLR), HCC transplant-OS (NLR), HCC transplant-DFS (NLR), Biliary Tract-OS (NLR), Lung-OS (NLR), Nasopharyngeal-OS (NLR), NSCLC-OS (NLR), Upper urinary and bladder-OS (NLR), Bladder-OS (NLR), CR prostate-OS (NLR), Renal-OS (NLR), Advanced renal-OS (NLR) |
| Suggestive (n=17) | p<10<sup>-4</sup>* with random effects;<br>>1000 individuals included | All cancers-OS (NLR), All cancers-OS (IN), Cervical-PFS (NLR), Ovarian-PFS (NLR), Gastric SR-OS (NLR), Colorectal-DFS (NLR), Colorectal-PFS (NLR), CLM SR-RFS (NLR), Colorectal SBR-DFS (NLR), HCC RFA-DFS (NLR), Nasopharyngeal-CSS (NLR), NSCLC-PFS (NLR), Urinary-OS (NLR), Upper urinary and bladder-CSS (NLR), Upper urinary and bladder-PFS (NLR), CR prostate-PFS (NLR), Renal-PFS (NLR), |
| Weak (n=29) | p<0.05* with random effects | All cancers-CSS (IN), Soft Tissue Sarcoma-OS (NLR), Gastric-DFS (NLR), Gastric-PFS (NLR), Biliary Tract-RFS (NLR), Pancreatic-CSS (NLR), Rectal-OS (NLR), Rectal-DFS (NLR), Rectal-RFS (NLR), CLM NS-OS (NLR), CLM NS-RFS (NLR), Colorectal PC-DFS (NLR), HCC and ICC-OS (IN), HCC mixed Tx-OS (NLR), HCC SR-OS (NLR), HCC RFA-OS (NLR), HCC TACE-OS (NLR), Lung-PFS (NLR), MPM-OS (NLR), NHC-OS (IN), Upper urinary-RFS (NLR), Bladder-RFS (NLR), Prostate-PFS (NLR), Urothelial carcinoma-OS (NLR), Urothelial carcinoma-RFS (NLR), Renal-OS (IN), Renal-RFS (NLR), Advanced renal-PFS (NLR), Localised renal-RFS (NLR) |
| Not Significant (n=9) | Not significant at p<0.05* with random effects | All cancers-OS (PN), All cancers-OS (SN), Gastric-OS (IN), Oesophageal-PFS (NLR), HCC SR-DFS (NLR), NSCLC-OS (IN), Localised prostate-OS (NLR), Renal-CSS (NLR), Localised renal-OS (NLR) |

(NLR) Neutrophil to lymphocyte ratio, (IN) Intratumoural neutrophils, (PN) Peritumoural neutrophils, (SN) Stromal neutrophils, (OS) Overall survival, (DFS) Disease-free survival, (PFS) Progression-free survival, (RFS) Reoccurrence-free survival, (CSS) Cancer specific survival, (PC) Palliative chemotherapy, (SR) Surgical resection, (CLM) Colorectal liver metastasis, (NS) Non-surgical, (HCC) Hepatocellular carcinoma, (ICC) Intrahepatic cholangiocarcinoma, (RFA) Radiofrequency ablation, (TACE) Trans-arterial chemoembolization, (MPM) Malignant pleural mesothelioma, (NSCLC) Non-small cell lung cancer, (NHC), Neck and head cancer, (CR) Castration resistant. \*p-values of random effects model. All cancers are defined as a grouping of cancer diagnosis unrelated to site, stage or treatment. No meta-analyses presented evidence of elevated neutrophils and improved cancer prognosis (HR<1).

Strong evidence was evident in three meta-analyses (4%) for associations between NLR and poor cancer prognosis. These associations included colorectal liver metastasis (CLM) (n=2) and nasopharyngeal cancer (n=1), with OS being the most frequently assessed outcome (n=2, 67%). Increased NLR was associated with reduced PFS in nasopharyngeal cancer and reduced OS in CLM, with and without surgical resection.

Twenty-three meta-analyses (28%) presented associations supported by highly suggestive evidence, including associations between increased NLR and poor prognosis in all cancers, breast, cervical, gynaecologic, gastric, biliary, pancreatic, colorectal, CLM, hepatocellular, lung, nasopharyngeal, non-small cell lung cancer (NSCLC), bladder, upper urinary and bladder, castration resistant prostate, renal and advanced renal cancer. The most commonly assessed outcome for highly suggestive associations was OS (n=17), followed by DFS (n=4), PFS (n=1) and RFS (n=1).

Seventeen meta-analyses provided suggestive evidence (21%) for an association between high NLR (n=16) or intratumoural neutrophils (n=1) and poor cancer prognosis, and twenty-nine meta-analyses provided weak evidence (36%) for an association between high NLR or intratumoural neutrophils and poor cancer prognosis. The association between intratumoural neutrophils and overall survival in all cancers was classified as suggestive, but there was weak evidence supporting associations with peritumoural neutrophils or stromal neutrophils. Details of the grading for each meta-analysis are included in *Appendix 7*.

### Quality assessment

The 26 meta-analyses categorised as providing either high suggestive (n=23) or strong evidence (n=3) arose from 20 individually published studies. Of these studies, three were ranked as critically low quality (15%), four as low quality (20%) and thirteen as moderate quality (65%). None of the assessed studies were ranked as high quality. The two studies which yielded the three meta-analyses categorised as providing strong evidence were both ranked as moderate quality (*Appendix 8*).

## DISCUSSION

A total of 81 associations between elevated NLR or TAN and cancer outcomes in 27 cancer sites were reviewed to assess the strength of the evidence supporting them. Three associations were supported by strong evidence and the evidence supporting associations including NLR was stronger than those including TAN. Although the studies included showed consistency in direction of effect, we detected poor reproducibility of findings and evidence of heterogeneity.

### Risk of elevated neutrophil to lymphocyte ratio

Previous studies have documented the prognostic role of neutrophils, particularly the NLR, and their link with poor outcomes for many cancer sites(13). The findings of this umbrella review support the association between elevated NLR and poor cancer prognosis, with 89% of the included meta-analyses providing a significant HR through random effects estimates (p<0.05) and no meta-analyses indicating a HR in the opposite direction of effect (HR<1).

All associations supported by strong evidence assessed elevated NLR in gastrointestinal and nasopharyngeal cancers. Nasopharyngeal cancers have also been linked to inflammation caused by smoking(82), presenting a potential confounder for the association. CLM represents a unique case where metastasis has already occurred and may present a link between elevated NLR and poor prognosis in metastasised cancers. Future research should assess the association between the NLR and prognosis in gastrointestinal cancers, with and without metastasis, and in respiratory cancers, by smoking status, to ensure these do not confound associations.

### Risk of intratumoural neutrophils

Although previous studies suggest a link between intratumoural neutrophils and the progression of cancer in the tumour microenvironment(15,17,83), the evidence for the association between intratumoural neutrophils and OS in all cancers was only classified as suggestive, with all other associations including TAN classified as weak or not significant. The significance of these associations may have been limited by small sample size and it is important to note that all meta-analyses considering TAN arose from a single study and may be to the same limitations.

It is important to note that all meta-analyses on associations concerning TAN arose from the same publication by Shen at al. 2014(48) and may be subject to the same limitations. The association between intratumoural neutrophils and cancer outcomes holds potential for a strong association due to the large effect size observed in this study and the plausibility of the biological mechanism behind the relationship(10). However, a recently published individual study on this association found that high levels of TAN had a protective effect in cancer(84), indicating that additional research is required to clarify the association.

### Risk of peritumoural and stromal neutrophils

Peritumoural and stromal neutrophils are hypothesised to play a role in creating a tumourigenic microenvironment through promotion of cellular proliferation and metastasis(17,85,86). However, the association for both peritumoural and stromal neutrophils and OS in all cancers was not supported by statistically significant evidence (p>0.05). Additional research is required to determine if any statistically significant association exists.

### Strengths

A key strength of this review comes from the use of umbrella review methodology, which only includes meta-analyses as evidence for quantitative data analyses(18). The use of meta-analyses ensures that effect size estimates are a balanced representation of the available evidence, as demonstrated by the sensitivity analysis of the association between elevated NLR and OS in rectal cancer from Dong et al. 2016(59) (*Appendix 9*). When an extreme outlier detected in this meta-analysis was removed from the analysis, the random effects estimate was not considerably altered due to the small weighting given to studies with large variances.

The TES did not detect a greater number of positive studies than expected in any meta-analyses, indicating that individual studies were not selectively included based on their significance(33). However, it is important to note that the TES is limited in power since biases that increase reporting of positive results will also increase the reported effect size, leading to a greater number of expected positive studies(33).

Although there are considerable differences between the included meta-analyses in terms of cancer site, stage and treatment, all the HR estimates reported in these meta-analyses were in the same direction of effect. This finding suggests consistency in the relationship between neutrophil indicators and poor outcomes across a wide spectrum of cancer diagnoses.

### Limitations

Only 34% of the identified meta-analyses were eligible for inclusion and 24% of the identified meta-analyses were excluded because they did not include sufficient data to be reproduced. Furthermore, we were unable to perfectly reproduce 26% of the included studies, highlighting issues with transparency and reproducibility of findings in epidemiologic research(87). Umbrella reviews also fail to include evidence published in individual studies after the last published meta-analyses. However, all the included meta-analyses in our study were recently published, with the oldest published in 2014, so the exclusion of individual studies in our case may be minimal. This exclusion of individual studies is of greatest concern in the association between TAN and cancer outcomes, due to a single meta-analysis being available.

The findings of this review are reliant on the quality of the included meta-analyses. This methodological limitation is of concern since 35% of the studies which yielded meta-analyses with highly suggestive or strong evidence were ranked as low or critically low quality by AMSTAR 2. There is also some concern over consistency, since meta-analyses aggregated the results of individual studies which categorised patients’ NLR or TAN levels as high or low using different cut off values (*Appendix 10*). Due to heterogeneity in these values, it is not possible to establish a dose-response relationship between neutrophil counts and cancer prognosis. Assessment of the individual studies included in each meta-analysis was outside of the scope of this review.

The assessment of heterogeneity using Cochrane’s Q test and the I^2^ statistic is problematic with varying study size. Cochrane’s Q test has weak power when there are few studies and excess power in detecting heterogeneity when studies are numerous, both of which are complications in this review(25).

### Causal association

This umbrella review does not address causality directly and cannot determine whether the association between neutrophils and poor prognosis in cancer is causal or due to confounding or reverse causality(88,89). Although previous studies have highlighted the paradoxical role of neutrophils in both tumour progression and suppression(17), this review suggests that the overall effect of high neutrophil counts is tumourigenic.

This review supports the relationship between elevated NLR and poor outcomes in cancer in terms of effect size and consistency of findings. We cannot address temporality as the studies included measured biomarkers before the initiation of treatment but generally after diagnosis. However, the biological mechanisms behind inflammation and cancer progression suggest temporality, as elevated NLR and TAN are proposed to promote increased cell proliferation(17), angiogenesis(90) and risk of metastasis(17,91) as contributors to poor prognosis.

### Clinical significance and future research

Future research should focus on strengthening the current evidence base for specific cancers which displayed suggestive and highly suggestive associations, addressing heterogeneity and, importantly, reverse causality. Unveiling a causal association between neutrophils and cancer outcomes could lead to cancer treatments which involve neutrophils as a therapeutic target and validate the NLR as a prognostic indicator in cancer. A casual association between neutrophils and poor prognosis could give further insight into experimental therapy which lowers neutrophils counts in the body to improve outcomes in cancer(85,92,93).

## CONCLUSION

The quantitative evidence presented suggests an association between elevated NLR and poor outcomes in cancer patients across a wide spectrum of diagnoses, stages of disease and courses of treatment. The evidence is strongest for associations between NLR and OS in CLM and nasopharyngeal cancer. The association between TAN and poor prognosis in cancer patients is limited by significant heterogeneity, small study effects and the existence of a single meta-analysis on the association. Further research is required to remove the limitations of the existing evidence.

## Acknowledgements

AJBT was supported by the Medical Research Council (UK MED-BIO Programme Fellowship, MR/L01632X/1). Funders had no role in data collection, analysis, interpretation or writing of the report. All authors had access to all the data in the study.

## Availability of data and materials

Data and computational code used for processing and analysis are available at https://github.com/megcupp/Neutrophils-and-Cancer-Prognosis.

All authors had full access to all of the data (including statistical reports and tables) in the study and can take responsibility for the integrity of the data and the accuracy of the data analysis.

## Authors’ contributions

AJBT conceived the study. MAC, AJBT and IT designed the study. MAC collected data and performed the analysis with input from MC, IT, ABJT and EE. MAC and AJBT wrote the manuscript with contributions from all authors. All authors critically revised and approved the manuscript. AJBT and MAC are the guarantors.

### Transparency declaration

The guarantors (AJBT and MAC) affirm that this manuscript is an honest, accurate, and transparent account of the study being reported; that no important aspects of the study have been omitted; and that any discrepancies from the study as planned (and, if relevant, registered) have been explained.

### Ethics approval and consent to participate

Not required.

## Supplementary Information

Supplementary Tables 1 - 2

Supplementary Figures 1 – 82

Appendix 1: Search strategy

Appendix 2: 239 extracted meta-analyses with exclusions identified

Appendix 3: Assessment of duplicate meta-analyses

Appendix 4: Description of meta-analyses with poor reproducibility

Appendix 5: Heterogeneity and small study effects in included meta-analyses

Appendix 6: Excess significance in included meta-analyses

Appendix 7: Grading of evidence

Appendix 8: AMSTAR Quality Rating of Meta-analyses with Highly Suggestive or Strong Evidence

Appendix 9: Sensitivity analyses

Appendix 10: NLR cut-offs of individual studies

Appendix 11: PRISMA 2009 Checklist

## REFERENCES

1. Siegel RL, Miller KD, Jemal A. Cancer statistics, 2017. CA Cancer J Clin. 2017;67(1):7–30.

2. Collaboration GB of DC. Global, regional, and national cancer incidence, mortality, years of life lost, years lived with disability, and disability-adjusted life-years for 32 cancer groups, 1990 to 2015: A systematic analysis for the global burden of disease study. JAMA Oncol. 2017 Apr;3(4):524–48.

3. Omran AR. The Epidemiologic Transition: A Theory of the Epidemiology of Population Change. Milbank Q. 2005;83(4):731–57.

4. Gospodarowicz M, O’Sullivan B. Prognostic factors in cancer. Semin Surg Oncol. 2003;21(1):13–8.

5. Allin KH, Nordestgaard BG. Elevated C-reactive protein in the diagnosis, prognosis, and cause of cancer. Crit Rev Clin Lab Sci. 2011 Aug;48(4):155–70.

6. Gupta D, Lis CG. Pretreatment serum albumin as a predictor of cancer survival: A systematic review of the epidemiological literature. Nutr J. 2010 Dec;9:69.

7. Danesh J, Collins R, Appleby P, Peto R. Association of fibrinogen, c-reactive protein, albumin, or leukocyte count with coronary heart disease: Meta-analyses of prospective studies. JAMA. 1998 May 13;279(18):1477–82.

8. Shankar A, Wang J, Rochtchina E, MC Y, Kefford R, Mitchell P. Association between circulating white blood cell count and cancer mortality: A population-based cohort study. Arch Intern Med. 2006 Jan 23;166(2):188–94.

9. Grimm R, Neaton J, Ludwig W. Prognostic importance of the white blood cell count for coronary, cancer, and all-cause mortality. JAMA. 1985 Oct 11;254(14):1932–7.

10. Powell DR, Huttenlocher A. Neutrophils in the Tumor Microenvironment. Trends Immunol. 2017 Aug;37(1):41–52.

11. Liang W, Ferrara N. The Complex Role of Neutrophils in Tumor Angiogenesis and Metastasis. Cancer Immunol Res. 2016;4(2):83–91.

12. Treffers LW, Hiemstra IH, Kuijpers TW, van den Berg TK, Matlung HL. Neutrophils in cancer. Immunol Rev. 2016;273(1):312–28.

13. Templeton AJ, McNamara MG, Seruga B, Vera-Badillo FE, Aneja P, Ocana A, et al. Prognostic role of neutrophil-to-lymphocyte ratio in solid tumors: a systematic review and meta-analysis. J Natl Cancer Inst. 2014;106(6):dju124.

14. Zahorec R. Ratio of neutrophil to lymphocyte counts—Rapid and simple parameter of systemic inflammation and stress in critically ill. Vol. 102, Bratislavské lekárske listy. 2001. 5–14 p.

15. Jensen HK, Donskov F, Marcussen N, Nordsmark M, Lundbeck F, von der Maase H. Presence of Intratumoral Neutrophils Is an Independent Prognostic Factor in Localized Renal Cell Carcinoma. J Clin Oncol. 2009 Oct;27(28):4709–17.

16. Granot Z, Henke E, Comen EA, King TA, Norton L, Benezra R. Tumor Entrained Neutrophils Inhibit Seeding in the Premetastatic Lung. Cancer Cell. 2017 Aug;20(3):300–14.

17. Coffelt SB, Wellenstein MD, de Visser KE. Neutrophils in cancer: neutral no more. Nat Rev Cancer. 2016 Jul;16(7):431–46.

18. Aromataris E, Fernandez R, Godfrey C, Holly C, Khalil H, Tungpunkom P. Summarizing systematic reviews: Methodological development, conduct and reporting of an umbrella review approach. Int J Evid Based Healthc. 2015 Sep;13:132–40.

19. Ioannidis JPA. Integration of evidence from multiple meta-analyses: a primer on umbrella reviews, treatment networks and multiple treatments meta-analyses. C Can Med Assoc J. 2009 Oct;181(8):488–93.

20. ProQuest. RefWorks [Internet]. Available from: https://refworks.proquest.com/

21. R Core Team. R: A language and environment for statistical computing [Internet]. R Foundation for Statistical Computing, Vienna, Austria. 2013. Available from: http://www.r-project.org/

22. Schwarzer G. Package “meta”: General Package for Meta-Analysis. Version 4.8-4 [Internet]. 2017. Available from: https://cran.r-project.org/web/packages/meta/meta.pdf

23. Riley RD, Higgins JPT, Deeks JJ. Interpretation of random effects meta-analyses. BMJ. 2011;342.

24. Hedges L V., Vevea JL. Fixed-and random-effects models in meta-analysis. Psychol Methods. 1998;3(4):486–504.

25. Higgins JPT, Thompson SG, Deeks JJ, Altman DG. Measuring inconsistency in meta-analyses. BMJ. 2003;327(7414):557–60.

26. Ioannidis JPA, Patsopoulos NA, Evangelou E. Uncertainty in Heterogeneity Estimates in Meta-Analyses. BMJ. 2007;335(7626):914–6.

27. Hoaglin DC. Misunderstandings about Q and “Cochran”s Q test’ in meta-analysis. Stat Med. 2016;35:485–95.

28. Salanti G, Ades AE, Ioannidis JPA. Graphical methods and numerical summaries for presenting results from multiple-treatment meta-analysis: an overview and tutorial. J Clin Epidemiol. 2011;64(2):163–71.

29. Gjerdevik M, Heuch I. Improving the error rates of the Begg and Mazumdar test for publication bias in fixed effects meta-analysis. BMC Med Res Methodol. 2014 Sep;14(1):109.

30. Egger M, Smith GD, Schneider M, Minder C. Bias in meta-analysis detected by a simple, graphical test. BMJ. 1997;315(7109):629–34.

31. Sterne JAC, Gavaghan D, Egger M. Publication and related bias in meta-analysis: Power of statistical tests and prevalence in the literature. J Clin Epidemiol. 2000;53(11):1119–29.

32. Sterne JAC, Sutton AJ, Ioannidis JPA, Terrin N, Jones DR, Lau J, et al. Recommendations for examining and interpreting funnel plot asymmetry in meta-analyses of randomised controlled trials. BMJ. 2011;343.

33. Ioannidis JPA, Trikalinos TA. An exploratory test for an excess of significant findings. Clin Trials. 2007 Jun;4(3):245–53.

34. Tsilidis KK, Papatheodorou SI, Evangelou E, Ioannidis JPA. Evaluation of Excess Statistical Significance in Meta-analyses of 98 Biomarker Associations with Cancer Risk. JNCI J Natl Cancer Inst. 2012;104(24):1867–78.

35. StataCorp. Stata Statistical Software: Release 14. College Station, TX: StataCorp LP; 2015.

36. StataCorp. power cox — Power analysis for the Cox proportional hazards model [Internet]. Available from: https://www.stata.com/manuals/psspowercox.pdf

37. R Core Team. Exact Binomial Test [Internet]. Statistical Data Analysis R. Available from: https://stat.ethz.ch/R-manual/R-devel/library/stats/html/binom.test.html

38. Papatheodorou SI, Tsilidis KK, Evangelou E, Ioannidis JPA. Application of credibility ceilings probes the robustness of meta-analyses of biomarkers and cancer risk. J Clin Epidemiol. 2015;68(2):163–74.

39. Salanti G, Ioannidis JPA. Synthesis of observational studies should consider credibility ceilings. J Clin Epidemiol. 2009;62(2):115–22.

40. Kyrgiou M, Kalliala I, Markozannes G, Gunter MJ, Paraskevaidis E, Gabra H, et al. Adiposity and cancer at major anatomical sites: umbrella review of the literature. BMJ. 2017 Feb;356.

41. Theodoratou E, Tzoulaki I, Zgaga L, Ioannidis JPA. Vitamin D and multiple health outcomes: umbrella review of systematic reviews and meta-analyses of observational studies and randomised trials. BMJ Br Med J. 2014 Apr;348.

42. Johnson VE. Revised standards for statistical evidence. Proc Natl Acad Sci. 2013 Nov;110(48):19313–7.

43. Thorlund K, Imberger G, Johnston BC, Walsh M, Awad T, Thabane L, et al. Evolution of Heterogeneity (I2) Estimates and Their 95% Confidence Intervals in Large Meta-Analyses. PLoS One. 2012 Jul;7(7):e39471.

44. Shea B, Reeves B, Wells G, Thuku M, Hamel C, Moran J, et al. AMSTAR 2: a critical appraisal tool for systematic reviews that include randomised or non-randomised studies of healthcare interventions, or both. BMJ. 2017;358:j4008.

45. Wickham H. Package ggplot2: Create Elegant Data Visualisations Using the Grammar of Graphics. 2016; Available from: https://cran.r-project.org/web/packages/ggplot2/index.html

46. Mei Z, Shi L, Wang B, Yang J, Xiao Z, Du P, et al. Prognostic role of pretreatment blood neutrophil-to-lymphocyte ratio in advanced cancer survivors: A systematic review and meta-analysis of 66 cohort studies. Cancer Treat Rev. 2017;58:1–13.

47. Paramanathan A, Saxena A, Morris DL. A systematic review and meta-analysis on the impact of pre-operative neutrophil lymphocyte ratio on long term outcomes after curative intent resection of solid tumours. Surg Oncol. 2014;23(1):31–9.

48. Shen M, Hu P, Donskov F, Wang G, Liu Q, Du J. Tumor-associated neutrophils as a new prognostic factor in cancer: a systematic review and meta-analysis. PLoS One. 2014;9(6):e98259.

49. Li Y, Liu X, Zhang J, Yao W. Prognostic role of elevated preoperative systemic inflammatory markers in localized soft tissue sarcoma. Cancer Biomarkers. 2016;(3):333–42.

50. Ethier J-L, Desautels D, Templeton A, Shah PS, Amir E. Prognostic role of neutrophil-to-lymphocyte ratio in breast cancer: a systematic review and meta-analysis. Breast Cancer Res. 2017;19(1):2.

51. Wu J, Chen M, Liang C, Su W. Prognostic value of the pretreatment neutrophil-to-lymphocyte ratio in cervical cancer: a meta-analysis and systematic review. Oncotarget. 2017;8(8):13400–12.

52. Ethier J-L, Desautels DN, Templeton AJ, Oza A, Amir E, Lheureux S. Is the neutrophil-to-lymphocyte ratio prognostic of survival outcomes in gynecologic cancers? A systematic review and meta-analysis. Gynecol Oncol. 2017;145(3):584–94.

53. Huanga Q, Zhou L, Zeng W, Qian-qian M, Wang W, Zhong M, et al. Prognostic Significance of Neutrophil-to-Lymphocyte Ratio in Ovarian Cancer?: A Systematic Review and Meta-Analysis of Observational Studies. Cell Physiol Biochem. 2017;41:2411–8.

54. Sun J, Chen X, Gao P, Song Y, Huang X, Yang Y, et al. Can the Neutrophil to Lymphocyte Ratio Be Used to Determine Gastric Cancer Treatment Outcomes? A Systematic Review and Meta-Analysis. Dis Markers. 2016;2016:7862469.

55. Zhang X, Zhang W, Feng L. Prognostic significance of neutrophil lymphocyte ratio in patients with gastric cancer: a meta-analysis. PLoS One. 2014;9(11):e111906.

56. Tang H, Lu W, Li B, Li C, Xu Y, Dong J. Prognostic significance of neutrophil-to-lymphocyte ratio in biliary tract cancers: a systematic review and meta-analysis. Oncotarget. 2017;8(22):36857–68.

57. Cheng H, Long F, Jaiswar M, Yang L, Wang C, Zhou Z. Prognostic role of the neutrophil-to-lymphocyte ratio in pancreatic cancer: a meta-analysis. Sci Rep [Internet]. 2015;5:11026.

58. Yang J-J, Hu Z-G, Shi W-X, Deng T, He S-Q, Yuan S-G. Prognostic significance of neutrophil to lymphocyte ratio in pancreatic cancer: a meta-analysis. World J Gastroenterol. 2015;21(9):2807–15.

59. Dong Y-W, Shi Y-Q, He L-W, Su P-Z. Prognostic significance of neutrophil-to-lymphocyte ratio in rectal cancer: a meta-analysis. Onco Targets Ther. 2016;9:3127–34.

60. Huang Y, Sun Y, Peng P, Zhu S, Sun W, Zhang P. Prognostic and clinicopathologic significance of neutrophil-to-lymphocyte ratio in esophageal squamous cell carcinoma: evidence from a meta-analysis. Onco Targets Ther. 2017 Feb;10:1165–72.

61. Zhang J, Zhang H-Y, Li J, Shao X-Y, Zhang C-X. The elevated NLR, PLR and PLT May predict the prognosis of patients with colorectal cancer: a systematic review and metaanalysis. Oncotarget. 2015;5(0).

62. Zheng D, Zheng C, Wu J, Ye H, Chen J, Zhou B, et al. Neutrophil-lymphocyte ratio predicts the prognosis of patients with colorectal cancer: A meta-analysis. Int J Clin Exp Med. 2016;9(1):78–90.

63. Tang H, Li B, Zhang A, Lu A, Xiang C. Prognostic significance of neutrophil-to-lymphocyte ratio in colorectal liver metastasis: A systematic review and meta-analysis. PLoS One. 2016;11(7):e0159447.

64. Malietzis G, Giacometti M, Kennedy RH, Athanasiou T, Aziz O, Jenkins JT. The emerging role of neutrophil to lymphocyte ratio in determining colorectal cancer treatment outcomes: a systematic review and meta-analysis. Ann Surg Oncol. 2014;21(12):3938–46.

65. Qi X, Li J, Deng H, Li H, Su C, Guo X. Neutrophil-to-lymphocyte ratio for the prognostic assessment of hepatocellular carcinoma: A systematic review and meta-analysis of observational studies. Oncotarget. 2016 Jul;7(29):45283–301.

66. Xue T-C, Zhang L, Xie X-Y, Ge N-L, Li L-X, Zhang B-H, et al. Prognostic significance of the neutrophil-to-lymphocyte ratio in primary liver cancer: a meta-analysis. PLoS One. 2014;9(5):e96072.

67. Xiao W-K, Chen D, Li S-Q, Fu S-J, Peng B-G, Liang L-J. Prognostic significance of neutrophil-lymphocyte ratio in hepatocellular carcinoma: a meta-analysis. BMC Cancer. 2014;14:117.

68. Sun X-D, Shi X-J, Chen Y-G, Wang C-L, Ma Q, Lv G-Y. Elevated Preoperative Neutrophil-Lymphocyte Ratio Is Associated with Poor Prognosis in Hepatocellular Carcinoma Patients Treated with Liver Transplantation: A Meta-Analysis. Gastroenterol Res Pract. 2016;

69. Zhao Q-T, Yang Y, Xu S, Zhang X-P, Wang H-E, Zhang H, et al. Prognostic role of neutrophil to lymphocyte ratio in lung cancers: a meta-analysis including 7,054 patients. Onco Targets Ther. 2015;8:2731–8.

70. Chen N, Liu S, Huang L, Li W, Yang W, Cong T, et al. Prognostic significance of neutrophil-to-lymphocyte ratio in patients with malignant pleural mesothelioma: a meta-analysis. Oncotarget. 2017;

71. Su L, Zhang M, Zhang W, Cai C, Hong J. Pretreatment hematologic markers as prognostic factors in patients with nasopharyngeal carcinoma: A systematic review and meta-analysis. Medicine (Baltimore). 2017;96(11):e6364.

72. Gu X-B, Tian T, Tian X-J, Zhang X-J. Prognostic significance of neutrophil-to-lymphocyte ratio in non-small cell lung cancer: a meta-analysis. Sci Rep. 2015;5:12493.

73. Wei Y, Jiang Y-Z, Qian W-H. Prognostic role of NLR in urinary cancers: a meta-analysis. PLoS One. 2014;9(3):e92079.

74. Luo Y, She D-L, Xiong H, Fu S-J, Yang L. Pretreatment Neutrophil to Lymphocyte Ratio as a Prognostic Predictor of Urologic Tumors: A Systematic Review and Meta-Analysis. Medicine (Baltimore). 2015;94(40):e1670.

75. Li X, Ma X, Tang L, Wang B, Chen L, Zhang F, et al. Prognostic value of neutrophil-to-lymphocyte ratio in urothelial carcinoma of the upper urinary tract and bladder: a systematic review and meta-analysis. Oncotarget. 2017;

76. Yin X, Xiao Y, Li F, Qi S, Yin Z, Gao J. Prognostic Role of Neutrophil-to-Lymphocyte Ratio in Prostate Cancer: A Systematic Review and Meta-analysis. Medicine (Baltimore). 2016;95(3):e2544.

77. Cao J, Zhu X, Zhao X, Li X-F, Xu R. Neutrophil-to-Lymphocyte Ratio Predicts PSA Response and Prognosis in Prostate Cancer: A Systematic Review and Meta-Analysis. PLoS One. 2016;11(7):e0158770.

78. Lucca I, Jichlinski P, Shariat SF, Rouprêt M, Rieken M, Kluth LA, et al. The Neutrophil-to-lymphocyte Ratio as a Prognostic Factor for Patients with Urothelial Carcinoma of the Bladder Following Radical Cystectomy: Validation and Meta-analysis. Eur Urol Focus. 2017 Aug;2(1):79–85.

79. Marchioni M, Cindolo L, Autorino R, Primiceri G, Arcaniolo D, De Sio M, et al. High Neutrophil-to-lymphocyte Ratio as Prognostic Factor in Patients Affected by Upper Tract Urothelial Cancer: A Systematic Review and Meta-analysis. Clin Genitourin Cancer. 2017;15(3):343–349.e1.

80. Na N, Yao J, Cheng C, Huang Z, Hong L, Li H, et al. Meta-analysis of the efficacy of the pretreatment neutrophil-to-lymphocyte ratio as a predictor of prognosis in renal carcinoma patients receiving tyrosine kinase inhibitors. Oncotarget. 2016;7(28):44039–46.

81. Boissier R, Campagna J, Branger N, Karsenty G, Lechevallier E. The prognostic value of the neutrophil-lymphocyte ratio in renal oncology: A review. Urol Oncol. 2017;35(4):135–41.

82. Zhu K, Levine RS, Brann EA, Gnepp DR, Baum MK. A population-based case-control study of the relationship between cigarette smoking and nasopharyngeal cancer (United States). Cancer Causes Control. 1995;6(6):507–12.

83. Galdiero MR, Bonavita E, Barajon I, Garlanda C, Mantovani A, Jaillon S. Tumor associated macrophages and neutrophils in cancer. Immunobiology. 2013 Nov;218(11):1402–10.

84. Berry RS, Xiong M, Greenbaum A, Mortaji P, Nofchissey A, Schultz F, et al. High levels of tumorassociated neutrophils are associated with improved overall survival in patients with stage II colorectal cancer. PLoS One. 2017;12(12):e0188799.

85. Gregory AD, McGarry Houghton A. Tumor-Associated Neutrophils: New Targets for Cancer Therapy. Cancer Res. 2011 Mar;71(7):2411 LP–2416.

86. Kuang D-M, Zhao Q, Wu Y, Peng C, Wang J, Xu Z, et al. Peritumoral neutrophils link inflammatory response to disease progression by fostering angiogenesis in hepatocellular carcinoma. J Hepatol. 2017 Aug;54(5):948–55.

87. Siegel V. Reproducibility in research. Dis Model Mech. 2011 May;4(3):279–80.

88. Höfler M. The Bradford Hill considerations on causality: a counterfactual perspective. Emerg Themes Epidemiol. 2005 Nov;2(1):11.

89. Fedak KM, Bernal A, Capshaw ZA, Gross S. Applying the Bradford Hill criteria in the 21st century: how data integration has changed causal inference in molecular epidemiology. Emerg Themes Epidemiol. 2015 Sep;12:14.

90. Jablonska J, Leschner S, Westphal K, Lienenklaus S, Weiss S. Neutrophils responsive to endogenous IFN-β regulate tumor angiogenesis and growth in a mouse tumor model. J Clin Invest. 2010 Apr;120(4):1151–64.

91. Coussens LM, Werb Z. Inflammation and cancer. Nature. 2002 Dec 19;420(6917):860–7.

92. Kim K, Skora AD, Li Z, Liu Q, Tam AJ, Blosser RL, et al. Eradication of metastatic mouse cancers resistant to immune checkpoint blockade by suppression of myeloid-derived cells. Proc Natl Acad Sci. 2014 Aug;111(32):11774–9.

93. Pekarek L, Starr B, Toledano A, Schreiber H. Inhibition of tumor growth by elimination of granulocytes. J Exp Med. 1995 Jan;181(1):435–40.

